# *easyfm*: An easy software suite for file manipulation of Next Generation Sequencing data on desktops

**DOI:** 10.1101/2021.09.29.462291

**Authors:** Hyungtaek Jung, Brendan Jeon, Daniel Ortiz-Barrientos

## Abstract

Storing and manipulating Next Generation Sequencing (NGS) file formats is an essential but difficult task in biological data analysis. The *easyfm* (**easy f**ile **m**anipulation) toolkit (https://github.com/TaekAndBrendan/easyfm) makes manipulating commonly used NGS files more accessible to biologists. It enables them to perform end-to-end reproducible data analyses using a free standalone desktop application (available on Windows, Mac and Linux). Unlike existing tools (e.g. Galaxy), the Graphical User Interface (GUI)-based *easyfm* is not dependent on any high-performance computing (HPC) system and can be operated without an internet connection. This specific benefit allow *easyfm* to seamlessly integrate visual and interactive representations of NGS files, supporting a wider scope of bioinformatics applications in the life sciences.

**Author summary:** The analysis and manipulation of NGS data for understanding biological phenomena is an increasingly important aspect in the life sciences. Yet, most methods for analysing, storing and manipulating NGS data require complex command-line tools in HPC or web-based servers and have not yet been implemented in comprehensive, easy-to-use software. This is a major hurdle preventing more general application in the field of NGS data analysis and file manipulation. Here we present *easyfm*, a free standalone Graphical User Interface (GUI) software with Python support that can be used to facilitate the rapid discovery of target sequences (or user’s interest) in NGS datasets for novice users. For user-friendliness and convenience, *easyfm* was developed with four work modules and a secondary GUI window (herein secondary window), covering different aspects of NGS data analysis (mainly focusing on FASTA files), including post-processing, filtering, format conversion, generating results, real-time log, and help. In combination with the executable tools (BLAST+ and BLAT) and Python, *easyfm* allows the user to set analysis parameters, select/extract regions of interest, examine the input and output results, and convert to a wide range of file formats. To help augment the functionality of existing web-based and command-line tools, *easyfm*, a self-contained program, comes with extensive documentation (hosted at https://github.com/TaekAndBrendan/easyfm) including a comprehensive step-by-step guide.

## 1 Introduction

With the broad implementation of NGS technologies in the life sciences, genomics and transcriptomics sequencing data are generated at an unprecedented rate [1–3]. Rapid progress in NGS technologies has brought massively high-throughput sequencing data to support research questions across many research fields, enabling a new era of genomic research [2,3]. Simultaneously, this advancement has brought enormous challenges in data analysis, of which efficient, standardized and consistent analysis are fundamental steps for maintaining reproducibility, especially for biologists [1,3]. However, many of the available tools for NGS data analysis require higher-order computational experience (e.g. various programming/scripting languages), expensive infrastructure (adequate HPC facilities and Cloud computing) and lack GUIs, making them inaccessible to many researchers, and cumbersome for even experienced biologists. Thus, the development of user-friendly standalone software for NGS data will accelerate the pace of research for scientists who have limited computer and bioinformatics experience.

NGS data processing often involves consecutive steps of trimming (including quality check), assembling, mapping, manipulating, converting and processing large files. FASTA [4] and FASTQ [5] file formats are generated by most NGS platforms, and further SAM/BAM [6], BED [7], GFF/GTF [8], and VCF [9] can be derived using FASTA and FASTQ files depending on the required analysis. The FASTA file, based on simple text, is the most basic format for reporting a sequence and is accepted by almost all sequence analysis programs. Each sequence starts with a “>” followed by the sequence name, a description of the sequence, and the sequence itself (nucleic acids or amino acids). The FASTQ file, a text-based format for storing both a biological sequence (usually nucleotide sequence) and its corresponding quality scores, is the most widely used format in sequence analysis and NGS sequencers. Each sequence requires at least 4 lines starting with “@” followed by the sequence, a “+” sequence identifier, and quality scores. Conveniently, FASTQ files can also be converted to FASTA files, the most commonly used file format for NGS data that enables direct sequencing of target genes. Many available tools (easySEARCH [10]; BlasterJS [11]; Sequenceserver [12]; orfipy [13]); Samtools and BCFtools [14] including *easyfm*) have not surprisingly focused on manipulating (analyse, collect, organise, interpret, and present data in meaningful ways) the FASTA file format to generate biologically relevant insights.

For the last decade, many HPC and Cloud-based NGS command-line programs or web-based platforms have wrapped popular high-level analysis and visualisation tools in an intuitive and appealing interface [15]. Galaxy (homepage: https://galaxyproject.org, main public server: https://usegalaxy.org, Australia: https://usegalaxy.org.au/) in particular has been successful in establishing itself as an analytics hub and an e-learning platform with global scientists, intending to produce accessible, reproducible and collaborative biological analyses [16,17]. Even with the huge achievements made in many analytical software packages and pipelines, further improvements in user-friendly standalone software are still required to facilitate the rapid discovery of meaningful sequences in very large data sets for novice users. To help augment the functionality of existing tools and allow for user-friendliness and convenience of NGS file manipulation, *easyfm* enables end-to-end file filtering, extracting and converting (FASTQ to FASTA) with a simple mouse click on desktops.

The *easyfm*, implemented in Python 3.7+, was developed with four work modules (Basic Local Alignment Search Tool [BLAST], BLAST-Like Alignment Tool [BLAT], Open Reading Frames [ORF], and File Manipulation) and a secondary window (Project Folder, Help and Log). Together, these modules and secondary window cover different aspects of NGS data analysis (mainly focusing on FASTA files), including post-processing, filtering, format conversion, and generating results. The functionality of each module has been described in the Results and Discussion section to have an easy-to-follow parallel comparison. *easyfm* is a GUI-based, lightweight but powerful, free and open-source desktop software for querying/manipulating NGS data sources and generating various outcomes. Since everyone can use it from anywhere to analyse data and find target sequences easily without any coding, HPC and/or internet/web-server connection, we hope the usefulness of *easyfm* can extend its potential use in a wide range of bioinformatics applications in the life sciences including teaching/learning materials in the classroom.

## 2 Design and implementation

*easyfm* can be used both by sophisticated data scientists and non-technical users who need an intuitive interface. The original intent for producing *easyfm* was to reduce reliance on any command lines/scripts or web-based platforms, by creating a standalone lightweight program with substantially reduced computational demands. *easyfm* provides key benefits in convenience, accessibility, and reproducibility because it does not include any heavyweight NGS data assembly, mapping and clustering workflows. *easyfm* can execute any pre-assembled genome/transcriptome FASTA files by selecting CPU numbers on a user’s desktop. While it mainly focuses on point-and-click analysis for less technical users, Log and Help functions could provide an interactive experience for monitoring and iterating on an executed code.

The *easyfm* work modules can provide support for post-processing, filtering, format conversion, and generating results to your given data (e.g. FASTA/Q files). It integrates four Python libraries and two executable programs with additional visualisation and conversation tools (mostly many well-established open-source Python packages) (Table 1). BLAST and indexing features provide the foundation for *easyfm* with approaches for all four work modules (BLAST, BLAT, ORF, and File Manipulation). While the user is required to select a module to execute, the user has full control over which input (including compressed files: *.gz) and output files/folders can be selected. *easyfm* also generates several output files (mostly in a tab-separated text file) that can be opened with standard text editors or Excel. To support work modules, *easyfm* also has a secondary window— Project Folder, Help and Log— that integrates with work modules (Fig 1). In addition, further assistance and information can be obtained via Help and Log to improve processes and performance. *easyfm* also contains all necessary dependencies. Simply unzip the folder and double-click easyfm.exe after downloading the program. Documentation, along with tutorials, is available at https://github.com/TaekAndBrendan/easyfm, and links to the *easyfm* download (https://github.com/TaekAndBrendan/easyfm/raw/main/windows/easyfm.7z).

**Fig 1.**
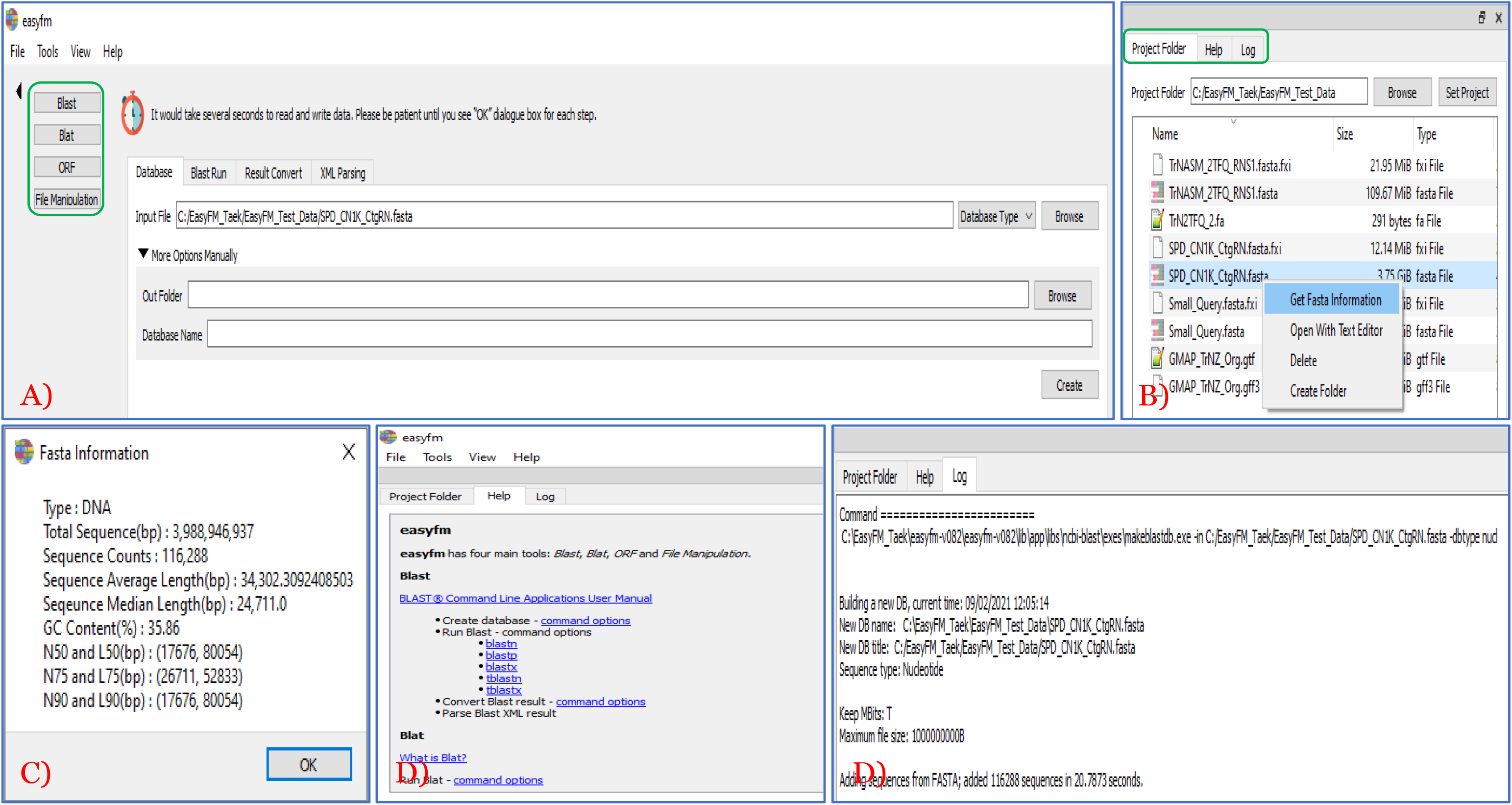
Integration of secondary window with main work module of *easyfm*. A) Four main work modules (green box) to BLAST, BLAT, ORF, and File Manipulation. B) Three secondary modules (green box) to assist with main work modules and extra features using a right mouse click. C) Fasta file stats information accessed from B. D) Adjustable secondary window (Help and Log) on the top and bottom.

**Table 1.** Software packages integrated into *easyfm* and their applications.

| Software | Application |
| --- | --- |
| Python Packages |  |
| Biopython [18] | Biopython is a set of freely available tools for common bioinformatics tasks including biological computation. |
| PyQt5 (https://pypi.org/project/PyQt5/) | PyQt is a Python binding of the cross-platform GUI toolkit Qt. |
| gffutils (https://github.com/daler/gffutils) | gffutils is for working with and manipulating the GFF and GTF format files typically used for genomic annotations. |
| Pyfastx [19] | The pyfastx is a lightweight Python C extension that enables users to randomly access sequences from plain and gzipped FASTA/Q files. |
| Executable Programs |  |
| BLAST+ (v2.11.0) [20] | BLAST+ is a sequence similarity searching tool with an enhancement of speed and query length. |
| BLAT (v3.2.1) [21] | BLAT is an alignment tool like BLAST and is useful for aligning long sequences and gapped mapping. |

Please note that *easyfm*, a self-contained program, includes all necessary dependencies and executable packages. Simply unzip the folder and double-click easyfm.exe after downloading the program.

## 3 Results and Discussion

### 3.1 Practical integration of secondary window in *easyfm*

To maximise the capability of the work modules (BLAST, BLAT, ORF, and File Manipulation), *easyfm* provides a secondary window, containing the tabs Project Folder, Help and Log to enhance the intuitive interface and interactive experience. As illustrated in Fig 1, the secondary window GUI components are freely adjustable with mouse movement (four corners) and are seamlessly integrated with the main work module of *easyfm*. The user can control input and output files by selecting the work folder (Project Folder and Set Project in Fig 1B green box) to use work modules. While the user can start with the default folder or select a specific work folder via Project Folder in the local drive, the input files must be in the designated folder. If the files (including compressed files: *.gz) are available in Project Folder, a simple right mouse click on the file can offer more options, such as Get Fasta Information (Stats), Open with Text Editor, Delete, and Create Folder (Fig 1B and 1C). The Help option is a resource intended to provide the end-user with information and support to *easyfm* work modules including its manual. To access additional information the user can click any of the links in Help (Fig 1D). Furthermore, to combine advanced functionalities with an easy-to-use interface, the Log option provides real-time log reporting and monitoring for every executed job (Fig 1D). This can aid in effective communication when reporting and resolving any program issues.

### 3.2 Intuitive interface of work modules in *easyfm*

To provide an integrated solution for NGS data file manipulation, *easyfm* provides an open-source tool with an easy installation and setup without relying on any web-based server or commercial licences. *easyfm* also allows users to consolidate the import/export data in FASTA/Q format (e.g. *.gz) under four work modules (BLAST, BLAT, ORF, File Manipulation) with an easy step-by-step process. *easyfm* is distributed under the MIT licence as all-in-one installer packages that contain all necessary software tools plus a manual explaining the analysis workflows step-by-step (https://github.com/TaekAndBrendan/easyfm).

#### 3.2.1 BLAST

BLAST is the most well-known analytics tool in life sciences and has become an essential program in every branch of biology to find regions of local similarity between biological (protein or nucleotide) sequences [18,22]. While the web-based National Center for Biotechnology Information (NCBI) BLAST suite of programs provides comprehensive sequence comparison, it is a major bottleneck due to delayed new data submission with embargo issues (including user-specific new data) and public availability on central BLAST repositories. Fortunately, BLAST can be installed and run locally, but its usage can be challenging for biologists who have limited experience of command-line interfaces. Furthermore, purchasing commercial software of a rich GUI-standalone tool (e.g. CLC Genomic Workbench and Geneious) and its licences is too expensive for many researchers and laboratories. To resolve these matters, *easyfm* provides a new Python-based free GUI for BLAST and more (Fig 2). Users can explore all BLAST+ (v2.11.0) features by creating a local database from which output format can be selected for including controlling analyses parameters and CPU cores. Even a common BLAST archive format (ASN.1) can be converted to any BLAST output format via Result Convert (Fig 2D). To save storage space and enable faster downstream analysis, a BLAST extensible markup language (XML) format (even generated externally) can be converted into a more compact form (e.g. a human-readable csv file) via XML Parsing (Fig 2E). Along with recent free tools [10–12], *easyfm* BLAST enables easy and seamless integration of visual and interactive representations of BLAST outputs supporting sequence similarity search. In particular, *easyfm* offers support for creating/searching a local database, changing format and parsing XML files as a standalone cross-platform application. This comprehensive and autonomous interface makes *easyfm* unique when compared to other free existing tools which need to rely on several different web servers.

**Fig 2.**
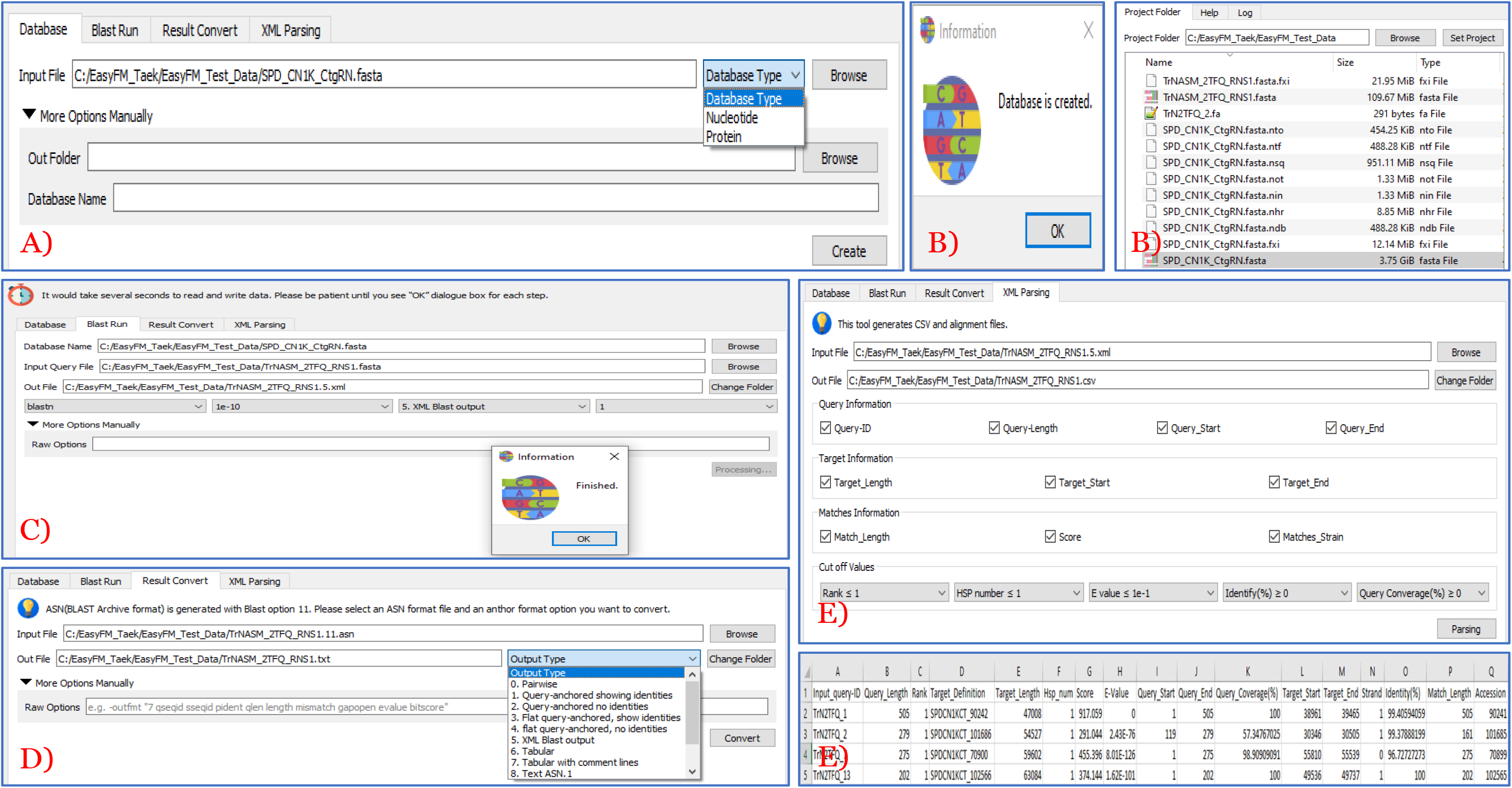
User-friendly standalone work modules in *easyfm*: BLAST module. Most steps include further manual options for a user-specified parameter. A) Create a local database by selecting nucleotide or protein. B) Job completion message and created database files listed in a secondary window. C) Run local BLAST with multiple features including output type. D) Convert from a BLAST archive file to a different output format. E) A BLAST xml file parsing with multiple options for a csv file.

#### 3.2.2 BLAT

BLAT is one of the alignment algorithms developed for the pairwise analysis and comparison of biological sequences with the primary goal of inferring homology to discover the biological function of genomic sequences [21]. While BLAT is less sensitive than BLAST, BLAT has a few clear advantages over BLAST from a practical standpoint in speed and convenience [23]. Compared to pre-existing pairwise sequence alignment tools, BLAT performed ∼500 times faster with mRNA/DNA alignments and ∼50 times faster with protein/protein alignments [21]. BLAT can be used either as a web-based server-client program (https://genome.ucsc.edu/cgi-bin/hgBlat) or as a standalone command-line program [23], but not a user-friendly GUI. However, *easyfm* BLAT (v3.2.1) enables users to control all parameters with a simple mouse click (Fig 3A) that can be a great advantage for novice biologists. Along with freely available *easyfm* BLAST, *easyfm* BLAT will simplify distributed computation pipelines to facilitate the rapid discovery of sequence similarities between NGS datasets. However, if the target genome and input sequences are big, using the standalone command-line BLAT in HPC is more suitable for batch runs, and more efficient than the web- and GUI-based BLAT because the standalone command-line in HPC can store more memory.

**Fig 3.**
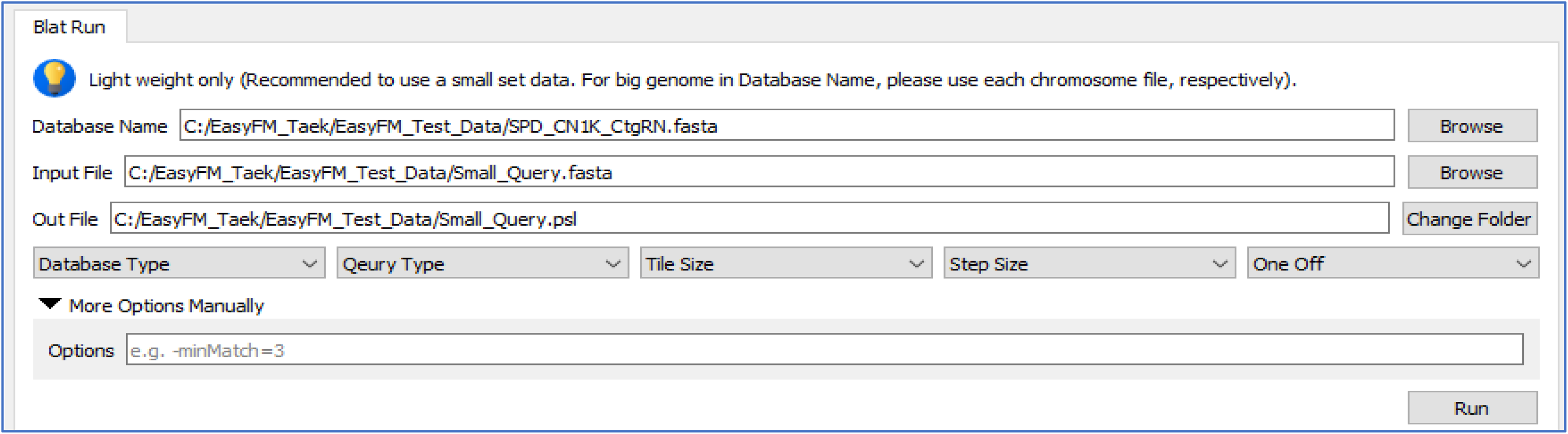
User-friendly standalone work modules in *easyfm*: BLAT module. Most steps include further manual options for a user-specified parameter. Create and run a local database with multiple options for a psl file that can open with text editor and Excel.

#### 3.2.3 ORF

An ORF(s) is the part of a reading frame that can be translated. The ORF (potential protein-coding sequence) is a continuous stretch of codons that usually begins with a start codon and ends at a stop codon. Understanding ORF(s) has become a piece of essential evidence to assist in gene prediction. As with other ORF finding tools, *easyfm* performs a six-frame translation of a nucleotide given a particular genetic code, finding all ORFs possible. Long ORFs are often used, along with other evidence, to initially identify candidate protein-coding regions or functional RNA-coding regions in a given DNA sequence, but the presence of an ORF does not necessarily mean that the region is always translated [24]. As BLAST and BLAT, the web-based ORF Finder (https://www.ncbi.nlm.nih.gov/orffinder/), ORF Predictor (http://bioinformatics.ysu.edu/tools/OrfPredictor.html) and command-line tools (ORF Investigator [25] and orfipy [13]) offer a range of ORF searches, but its usage can be challenging for biologists due to lack of computer programming literacy and limited query sequence length. To maximise the flexibility, the *easyfm* ORF provides a fast and efficient approach for all possible translation and extraction of ORFs from nucleotide sequences (FASTA format of nucleotide and protein output from six-frame translation) (Fig 4). With a simple mouse click solution, users can compare the translated outcomes with their biological evidence to avoid false discovery as well as control specific parameters without any limitation of query sequence length. Along with existing tools [13,25], *easyfm* ORF will provide rapid, flexible searches in multiple output formats to allow the easy downstream analysis of ORFs.

**Fig 4.**
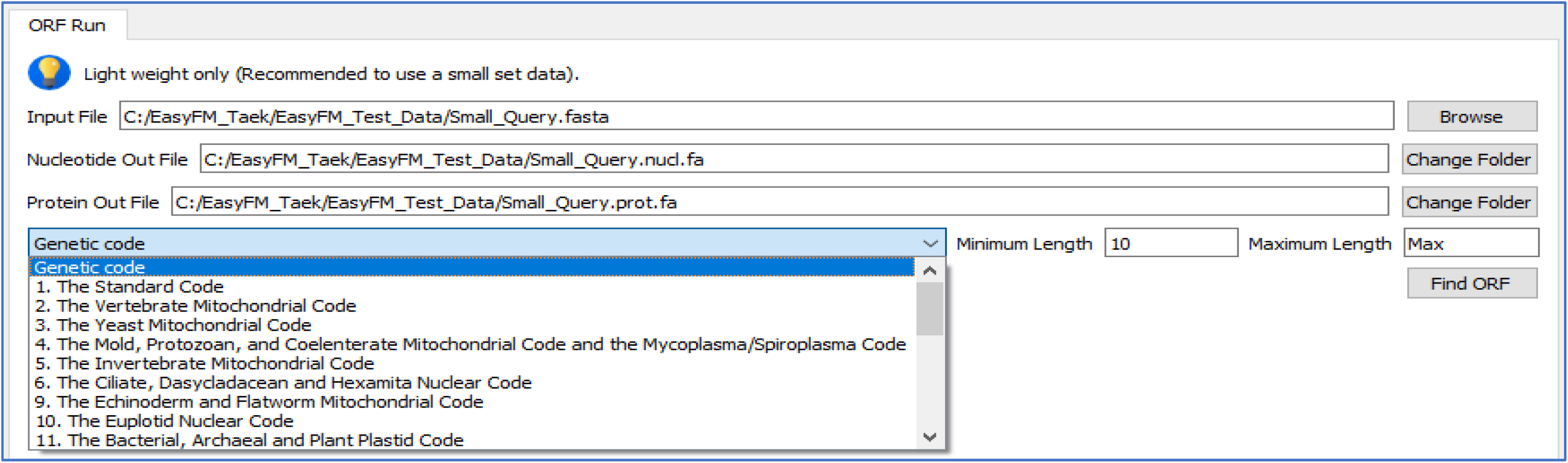
User-friendly standalone work modules in *easyfm*: ORF module. Most steps include further options for a user-specified parameter. Run ORF with different genetic codes for coding and protein sequences. A FASTA format output file of nucleotide and protein from a six-frame translation will be generated.

#### 3.2.4 File Manipulation

Various file formats have been introduced with the development of different DNA/RNA sequencing technologies. While there are many different biological file formats related to NGS analyses (or to store and manipulate), FASTA/Q files are most commonly encountered in the bioinformatics community. This is due to their flexibility: FASTA/Q files can be read, mapped and indexed by several different software packages to generate SAM/BAM, GFF/GTF, VCF, and more. Using a fai index file in conjunction with a FASTA/Q file containing reference sequences enables efficient access to arbitrary regions within those reference sequences and extracts subsequences from the indexed reference sequence (Danecek et al. 2021; Quinlan & Hall 2010).

Like other modules, the web-based Galaxy (homepage: https://galaxyproject.org, main public server: https://usegalaxy.org, Australia: https://usegalaxy.org.au/) and command-line tools (Samtools and BCFtools [14]; BEDTools [26]) offer a range of NGS data file manipulation capabilities, but its usage can be challenging for biologists due to lack of computer language literacy and internet dependence. To enhance and extend the flexibility and convenience, we present *easyfm*, a free single GUI for NGS file manipulation (mainly for FASTA files) (Fig 5). Since users can control everything with a simple mouse click on a desktop, the tools available in the *easyfm* would be a convenient way to teach bioinformatics/data analysis, and to quickly analyse results without being hampered by command line tools and HPC Secure Shell (SSH) connections.

**Fig 5.**
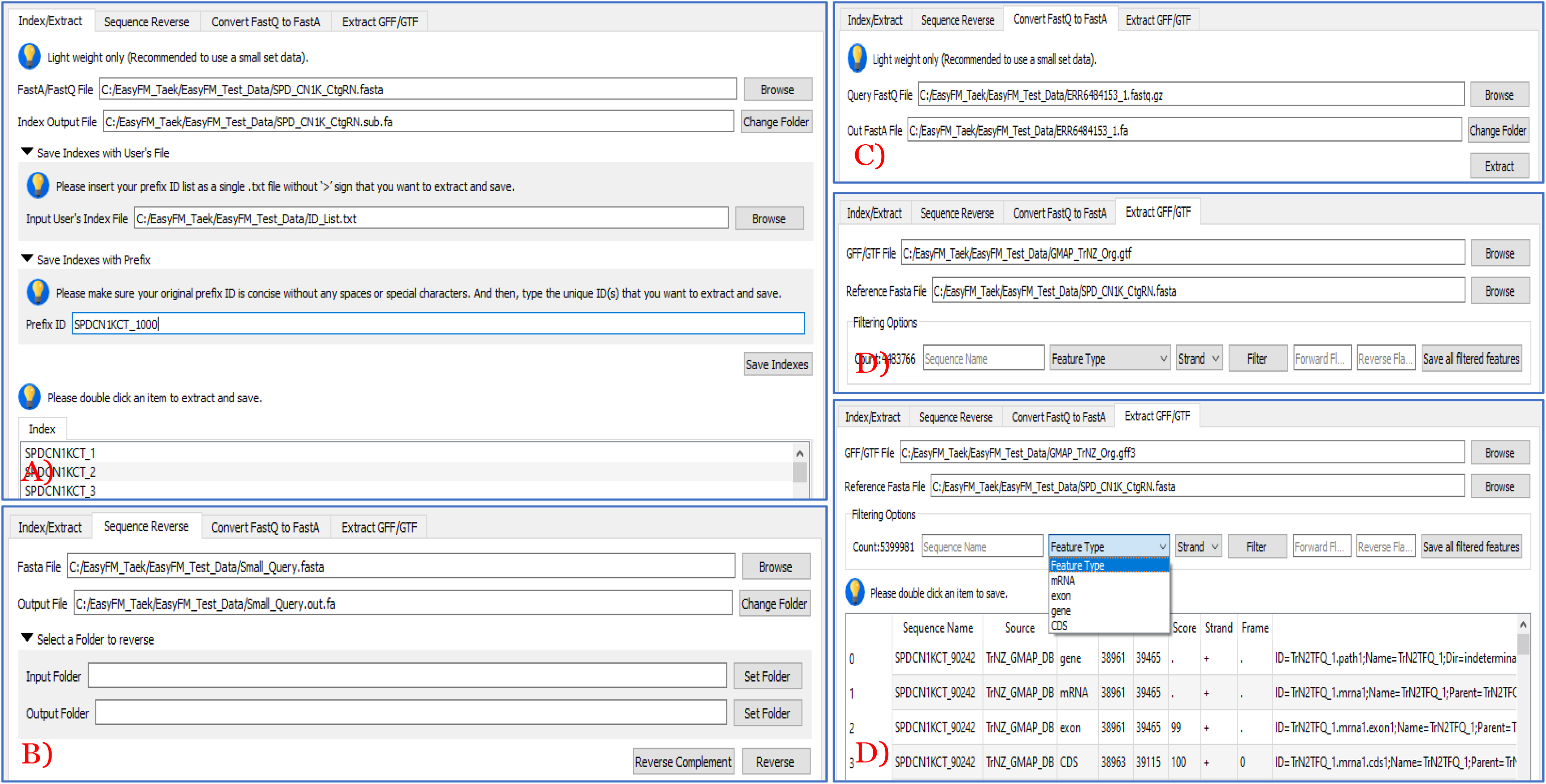
User-friendly standalone work modules in *easyfm*: File Manipulation module. Most steps include further individual selection by manually saving as a FASTA file for a user-specified sequence ID. A) Select a FASTA file to index. B) Convert nucleotide sequences for reverse complement or just reverse sequence. C) Convert and extract from FASTQ to FASTA. D) Extract sequences with specific IDs from indexed reference FASTA and GFF3/GTF files with different features.

Users can import any FASTA/Q files to index and extract the indexed ID with its sequence by double-clicking, matching Prefix ID and selecting a provided text file (Fig 5A). Even the FASTQ file can be converted to the FASTA file and the given FASTA file change its direction via Reverse Complement and Reverse (Fig 5B and 5C). For wide applications, *easyfm* File Manipulation also allows users to easily manipulate (including filtering [IDs, features and strand] and extracting sequence regions) and consolidate from GFF and GTF files if its corresponding reference genome/transcriptome sequences are present (Fig 5D). To enhance user-friendliness, users can extract a given sequence as a FASTA file with extra flanking regions for both directions by entering the desired sequence length (numeric numbers). Along with existing tools [14,26], *easyfm* File Manipulation will provide a stable and modular platform for manipulating sequence data and files to ensure high reproducibility standards in the NGS era.

## Availability and future directions

*easyfm* is implemented in Python and available under the MIT license and works on Windows, Linux and Mac systems. This package is also available on PyPI python package manager. The current code runs under Python 3.7+ and virtualenv. Other dependency includes gffutils, pyfastx, PyQt5 and Biopython (Table 1). More information and the manual may be obtained from the website: https://github.com/TaekAndBrendan/easyfm.

In the future, we will continue to update the toolbox with new fast and easy GUI support, including new embedding methods such as DIAMOND [27,28], (Buchfink et al. 2015, 2021) and pBLAT [29] with low resource requirements and both multithread and cluster computing support, making these methods suitable for running on standard desktops and laptops. Future versions of *easyfm* will also include additional integration points allowing us to intersect, merge, count, complement, and shuffle genomic intervals from multiple files in widely-used genomic file formats such as BAM, BED and VCF.

## Acknowledgements

The authors are grateful to their colleagues (special thanks to Maddie James, Melanie Wilkinson, Cara Conradsen, and Nicholas O’Brien) and collaborators for their valuable and constructive comments.

## Notes

### Competing Interest Statement

The authors have declared no competing interest.

